# A novel recombinant DNA system for high efficiency affinity purification of proteins in *Saccharomyces cerevisiae*

**DOI:** 10.1101/025205

**Authors:** Brian H. Carrick, Lixuan Hao, Philip J. Smaldino, David R. Engelke

## Abstract

Isolation of endogenous proteins from *Saccharomyces cerevisiae* has been facilitated by inserting encoding polypeptide affinity tags at the C-termini of chromosomal open reading frames (ORFs) using homologous recombination of DNA fragments. The tagged protein isolation is limited by a number of factors, including high cost of affinity resins for bulk isolation and low concentration of ligands on the resin surface, leading to low isolation efficiencies and trapping of contaminants. To address this we have created a recombinant “CelTag” DNA construct from which PCR fragments can be created to easily tag C-termini of *S. cerevisiae* ORFs using selection for a *nat1* marker. The tag has a C-terminal cellulose binding module to be used in the first affinity step. Microgranular cellulose is very inexpensive and has an effectively continuous ligand on its surface, allowing rapid, highly efficient purification with minimal background in a single step. Cellulose-bound proteins are released by specific cleavage of an included site for TEV protease, giving nearly pure product. The tag can be lifted from the recombinant DNA construct either with or without a 13x myc epitope tag between the target ORF and the TEV protease site. Binding of CelTag protein fusions to cellulose is stable to high salt, nonionic detergents, and 1 M urea, allowing stringent washing conditions to remove loosely associated components, as needed, before specific elution. It is anticipated that this reagent will allow isolation of rare or unstable protein complexes from large quantities of yeast extract, including soluble, membrane-bound, or chromatin-associated assemblies.

## INTRODUCTION

Proteomics requires quick, specific, and dependable methods for purifying proteins and complexes. Typically techniques for characterizing protein complexes require large quantities of highly purified protein, which is difficult to obtain for rare or unstable complexes. The most common method currently available for specific and pure protein purification in yeast is the dual affinity tag, known as TAP (tandem affinity purification) tag (Rigaut *et al.* 1999). This tag uses the IgG binding domain of protein A of *Staphylococcus aureus* (ProtA) (Uhlen *et al.* 1983; Nilsson *et al.* 1987) and a calmodulin-binding peptide separated by a Tobacco Etch Virus (TEV) protease cleavage site (Dougherty *et al.* 1989; Kapust *et al.* 2001). The tag is added on to the C-terminal end of proteins, which is bound to IgG immobilized on a bead matrix during the first step of purification. The bound protein is eluted from the matrix by TEV protease cleavage under mild conditions, and then is bound to calmodulin-coated beads in the presence of calcium during the second purification step. The bound proteins are then eluted from the calmodulin beads with EGTA (Lohman *et al.* 1989; Puig *et al.* 2001; Van Driessche *et al.* 2005).

Tandem affinity methods are very powerful, yet have limitations. Native protein complexes are often rare and unstable, and therefore without rapid, high yield purification methods these complexes might not be isolated cleanly using standard purification techniques. Often *in vivo* overexpression of rare protein complexes is impractical because of artificial interactions and low concentrations of physiologically relevant partners. Such complexes, whether soluble, bound to specific genes, or associated with membranes, can exist in less than one copy per cell, making isolation on a large scale mandatory. The problems with current approaches are that affinity purification reagents are expensive on a large scale, and the low concentration of affinity ligand per surface area/volume on affinity resins makes specific binding inefficient, leading to high signal-to-noise ratios. We sought to create a new recombinant DNA system for adding affinity tags to proteins that provides enough specificity and enrichment in the purification at relatively low cost.

The result is an affinity tag that provides enough specificity and enrichment in the purification while also maintaining a relatively low cost, especially during the first, bulk isolation step. To do this we created a C-terminal dual tag (termed CelTag) consisting of a family 3 cellulose binding module (CBM3) (Levy and Shoseyov 2002) and a 13x c-Myc repeat epitope tag (Terpe 2003) separated by a TEV protease cleavage site (Dougherty *et al.* 1989) (Figure 1). CBM3 was chosen due to its high affinity and specificity for a cellulose matrix and it has been shown to form an independent domain at either terminus of a protein (Stofko-Hahn *et al.* 1992; Levy and Shoseyov 2002; Hong *et al.* 2007). Cellulose is an excellent matrix for purification due to its very low cost, low background due to the few proteins that have nonspecific affinity for it (in yeast and animals), and its stability in a number of buffer conditions and pH levels. Recently it was shown that CBM3 could be used as a single affinity tag to purify overexpressed proteins from recombinant DNA in *Pichia pastoris* (Wang and Hong 2014). A 13x c-myc repeat epitope was added in our constructs due to the availability and relatively low cost of strong monoclonal antibodies produced against this sequence. This could be particularly useful for lesser studied proteins in which only poor or no antibodies exist to study them.

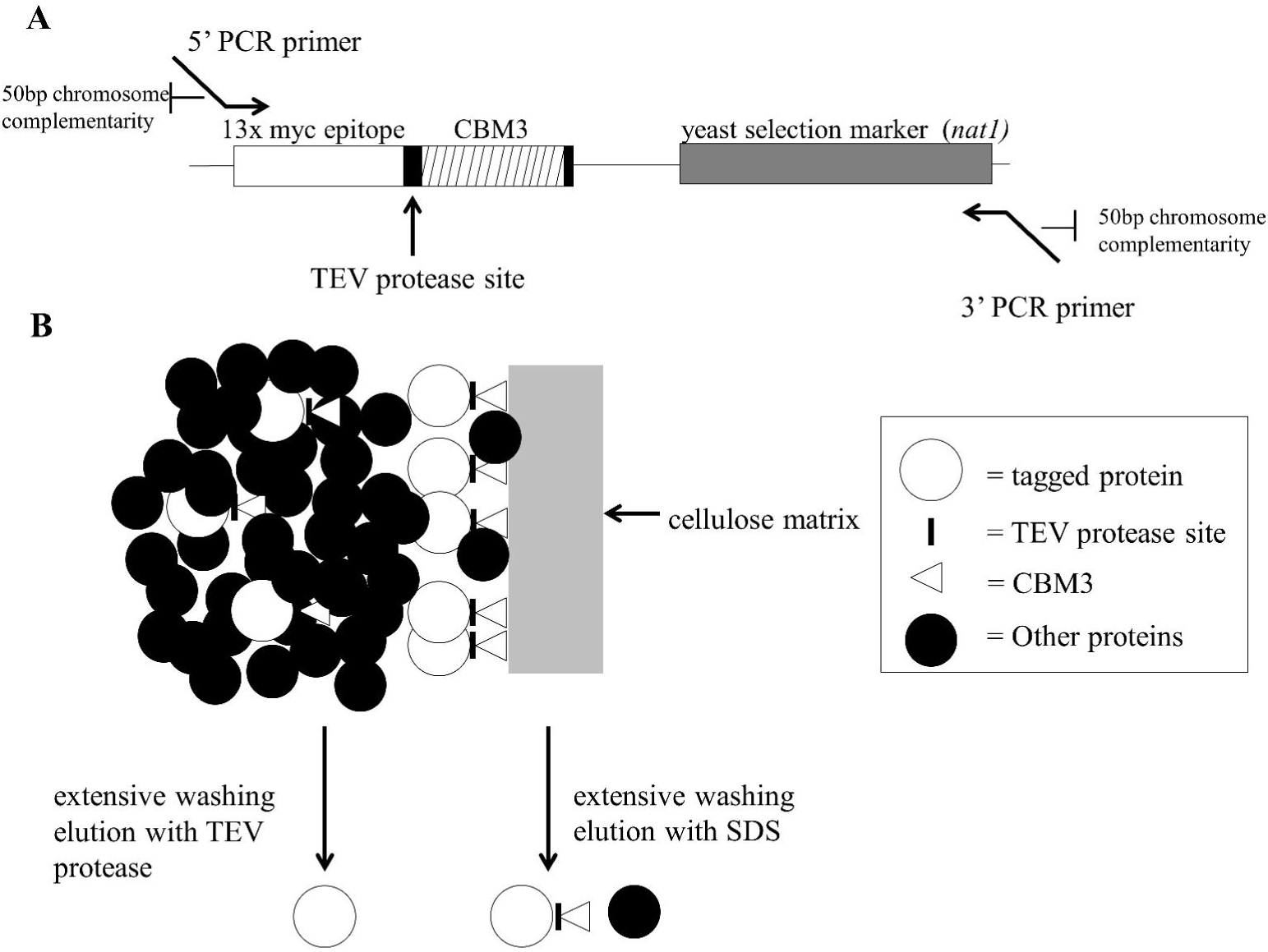

A major advantage of yeast for this type of study is that the transformed DNA sequences recombine into the yeast chromosomes by sequence homology (Figure 1) at a relatively high frequency. This makes it possible to easily tag the C-termini of proteins and analyze them *in vivo* in the proper physiological context. To determine the effectiveness and efficiency of integration, a marker is engineered into the tag, which allows for the yeast transformants to be isolated by growth on selective media. Here we provide a proof of concept example of isolation of a soluble yeast protein

## MATERIALS AND METHODS

All reagents and chemicals are reagent grade and purchased from Sigma-Aldrich (St. Louis, MO, USA) and Fisher Scientific (Pittsburgh, PA, USA) unless otherwise noted.

### Strains

*E. coli* DH5α strain (F– Φ80*lac*ZΔM15 Δ(*lac*ZYA-*arg*F) U169 *rec*A1 *end*A1 *hsd*R17 (rK–, mK+) *pho*A *sup*E44 λ– *thi*-1 *gyr*A96 *rel*A1) was used as the host cell for plasmid manipulation. Luria-Bertani (LB) medium with or without 100 μg/mL ampicillin was used to culture *E. coli* host cell or transformants. *S. cerevisiae* BY4741 (MATα *his*3Δ1, *leu*2Δ, *met*15Δ, *ura*3Δ) was used as a parent strain for transformation with DNA fragments as well as for recombinant protein expression. YPD medium (1% yeast extract, 2% peptone, and 2% dextrose) with or without 150 μg/mL of clonNat (Werner BioAgents, Meisenweg, Jena, Germany) was used to culture *S. cerevisiae* parent strains and transformants

### Plasmid Preparation

All plasmids constructed in this study were isolated in DH5α cells through QIAGEN mini or midi prep according to procedures outlined in the QIAGEN Plasmid Purification Handbook (http://www.qiagen.com/literature/render.aspx?id=369).

### Construction of CelTag Plasmid

A PCR fragment containing the 13x myc epitope repeat and clonNAT resistance gene was created from the digestion of the pFA6a-13myc-natMX6 plasmid (gift from Dr. Antony Carr at University of Sussex) (Van Driessche *et al.* 2005) with restriction enzymes Not1 and Cla1(New England Biolabs, Ipswich, MA) at 37°C overnight. The fragment was then ligated into the pRS426 (Sikorski and Hieter 1989) which was cut with the same enzymes. A fragment containing the CBM3 was obtained from the pCIG plasmid (gift from Dr. Jiong Hong at Virginia Polytechnic Institute and State University) (Hong *et al.* 2008) and gap repaired between the 13x myc repeat and clonNAT resistance gene (See Supplemental File S1 for complete sequence). To avoid carrying over a yeast-compatible expression plasmid when transforming yeast with amplified DNA fragments, the tag was moved from pRS426 (shuttle vector for yeast and bacteria) to the pGEM-T Easy vector (Promega, Madison, WI) (a bacterial only plasmid). A DNA fragment containing the entire tag and selection marker was PCR-amplified from the pRS426 vector by PCR and was ligated into the pGEM-T Easy vector according to manufacturer’s specifications (https://www.promega.com/resources/protocols/technical-manuals/0/pgem-t-and-pgem-t-easy-vector-systems-protocol/). Correct sequence of the tag was confirmed by Sanger sequencing (University of Michigan Sequencing Core). Plasmid will be available through Addgene (Plasmid #66562) and the sequence and full map are provided in Supplementary Figure 1.

### DNA Fragment Purification

Fragments of interest for construction of the CelTag and for the creation of tagged ORFs were isolated by running approximately 25 μg of DNA on a 0.8% agarose electrophoresis gel containing 20 mg/mL of ethidium bromide for visualization of the DNA. Gel bands of interest were quickly excised under long wave UV-light and DNA was electroeluted from the gel piece by sealing it in Spectra/Por Dialysis Membrane (MWCO 12000-1400) (Spectrum Labs, Rancho Dominguez, CA) and placed in 0.5X TBE buffer at 150V until no ethidium bromide was visible in gel slices. The DNA was then extracted using phenol/chloroform and precipitated with NaOAc and EtOH.

### PCR of CelTag Fragment

Primers are designed as follows; the forward primer contains 50 base pairs of homology to the end of the open reading frame where the fragment is to be inserted (excluding the stop codon), followed by 25 base pairs of complementarity to the beginning of the CelTag recombinant DNA construct (Figure 1). The reverse primer is 50 base pairs of homology to sequences downstream of the target ORF (∼25 nucleotides downstream from the stop codon), with 25 nucleotides complementary to the end of the CelTag fragment (See Supplementary Figure S2 and Table S1for sequences). The tag was amplified from the plasmid with the homologous ends to the ORF of interest and was purified by electrophoretic DNA purification and extraction as described above. To ensure there was enough fragment for efficient transformation into yeast, a total of 400 μL of PCR was done (4 × 100 μL reactions).

### PCR Conditions

Optimum PCR conditions vary slightly with different primers, but were approximately as follows. Reaction consisted of Pfu Buffer (1X: 10 mM KCl, 6 mM (NH_4_)_2_SO_4_, 2 mM MgCl_2_, 20 mM Tris-HCl pH 8.8, 0.1% Triton X-100, and 0.1 mg/mL BSA), dNTPs (100μM each), primers (1 μM each), Taq polymerase (Engelke *et al.* 1990) (5 units), Pfu DNA polymerase (Lu and Erickson 1997) (1 unit). Cycles: (94°C, 5:00) x1 cycle; (94°C, 30 seconds), (48°C, 30 seconds), (72°C, 3 minutes) x5 cycles; (94°C, 30 seconds), (53°C, 30 seconds), (72°C, 3 minutes) × 30 cycles; (72°C, 7 minutes), (4°C, hold).

### Transformations

Saturated 5 mL overnight cultures of BY4741 grown in YPD were used to seed 22 mL cultures of YPD to an OD_600nm_ of approximately 0.2. Cultures were grown at 30°C until the OD_600nm_ was 0.6-0.8 (mid-log phase). 10 mL of cells were aliquoted for each transformation and were spun at 3500 RPM (Sorvall T 6000 B) for 5 minutes and washed twice with 100 mM lithium acetate (LiOAc). Pellets were then resuspended in 100 μL of 100 mM LiOAc. 10 μg of salmon sperm DNA and approximately 25 μg of purified PCR fragment containing the CelTag fragment with ends homologous to chromosomal sequences, were added to the transformation reaction and incubated at 30°C for 15 minutes. Six hundred μL of PEG Mix (100 mM LiOAc, 50% PEG 3350) was added and mixed well before incubation at 30°C for 30 minutes. Then, 68 μL of DMSO was added and then reactions were heat shocked at 42°C for 15 minutes. Cells were spun at 3500 RPM for 5 minutes and resuspended in 1 mL of YPD media and then grown at 30°C for 4 hours. The cells were then spun at 3500 RPM for 5 minutes and resuspended in 250 μL of dH_2_O before plating to YPD+1.5XclonNAT (150 μg/mL clonNAT). Positive clones were screened for by PCR over insertion junctions as well as visualization of myc-epitope tagged protein by western blot analysis using an anti-Myc-repeat antibody (9E10) (Santa Cruz Biotech).

### Yeast Cell Lysate Preparation

Control or transformed yeast cells in 6 L of YPD medium were grown to early log phase, an OD_600nm_ of approximately 0.2 (either wild-type BY4741 or with tag inserted and therefore clonNAT resistant). Cells were harvested at 4000 RPM for 5 minutes at 4°C (Sorvall RC 5C Plus centrifuge using SLA-3000 rotor), resuspended in 300 mL of Lysis Buffer 1 (1 M sorbitol, 50 mM Tris-HCl pH 7.2, 10 mM DTT), recovered by centrifugation at 4°C and resuspended in 120 mL of Lysis Buffer 2 (1 M sorbitol, 50 mM Tris-HCl pH 7.2, 3 mM DTT, 0.4 mg/mL Zymolyase 20T). The cell suspension was incubated at 30°C for 30 minutes with gentle swirling. Cells were spun at 4000 RPM for 5 minutes at 4°C and washed gently, but thoroughly with 120 mL Lysis Buffer 3 (1 M sorbitol, 10 mM Tris-HCl pH 7.2). The cells were harvested at 4000 RPM for 5 minutes at 4°C (using the same rotor as above), followed by resuspension in 60 mL CBM3 Binding Buffer (50 mM Tris-HCl pH 6.5, 100 mM NaCl, 1 mM EDTA (Wan *et al.* 2011) containing cOmplete, Mini, EDTA-free Protease Inhibitor Cocktail (Roche, Basel, Switzerland). The suspension was sonicated (Branson Digital Sonifier 450 using the 102-C converter, ^1/2^” tapped bio horn, and 1/8” tapered microtip as the probe assembly) on ice for a total of 1.5 minutes (3 seconds on, 10 seconds off) at 25% amplitude. Debris was spun out at 4000 RPM for 5 minutes at 4°C and lysate was then aliquoted for binding experiments and stored at - 80°C.

### Preparation of Regenerated Amorphous Cellulose (RAC)

RAC was prepared by acid treatment as previously described (Wang and Hong 2014). The resulting material is a rather gelatinous and can be difficult to work with. After the acid treatment, the material was resuspended once in CBM3 binding buffer with 2M salt, once in CBM3 binding buffer with no salt, and 3 times in CBM3 binding buffer. The RAC was resuspended and stored in a 1:1 slurry of cellulose and binding buffer.

### Preparation of Microgranular Cellulose

Cellulose was washed 3 times in an 20 volume excess of 2 M NaCl CBM3 binding buffer, 2 times in no salt CBM3 binding buffer, and 3 times in CBM3 binding buffer. Cellulose was resuspended in a 1:1 slurry with CBM3 binding buffer.

### Cellulose Pulldowns

For each pulldown reaction, 450 μL of cell lysate as prepared above, was used to resuspend variable volumes of packed cellulose pellet and allowed to bind at 4°C for 20 minutes with gentle agitation. Cellulose pellets were washed four times in an excess volume (50-fold) of binding buffer before elution in 50 μL of SDS-Page Loading Buffer (1X working: 80 mM Tris-HCl pH 6.8, 10% glycerol, 2% SDS, 100mM DTT, 0.2% bromophenol blue, 100 mM 2-Mercaptoethanol), a binding buffer with varied additives, or a TEV protease elution.

### TEV Protease Elution

TEV protease was purified using a previously described plasmid encoding a His6 tagged TEV clone and published protocol (Tropea *et al.* 2009). TEV elutions were done in 100 μL of TEV protease buffer (50 mM Tris-HCl pH 8.0, 0.5 mM EDTA, 1 mM DTT) containing varying amounts of TEV protease at 30°C for 1 hour. Units of TEV were determined by comparison to AcTEV protease (Life Technologies, Carlsbad, CA). TEV protease was removed by the PureProteome Nickel Magnetic Bead System (EMD Millipore, Billerica, MA) using manufacturer’s instructions. (http://www.emdmillipore.com/US/en/product/PureProteome%E2%84%A2-Nickel-Magnetic-Bead-System,MM_NF-LSKMAGH02?bd=1#documentation)

### Protein Visualization and Quantification

For western blot analysis, SDS-PAGE loading buffer was added to each sample and heated at 95°C for 5 minutes, spun briefly in a microcentrifuge to remove debris, and subjected to electrophoresis on 4-15% gradient, 15 well SDS-Page gels (BioRad, Hercules, CA). The general western blot protocol at http://www.abcam.com/protocols/general-western-blot-protocol was used for all western blots. PVDF membrane (Millipore, Billerica, MAwas used for all transfers. Primary antibody incubation was done with a 1:400 dilution of an anti-Myc-repeat antibody (9E10) (Santa Cruz Biotech, Dallas, TX) for 1 hour at 4°C, and a secondary antibody incubation was performed with a 1:1000 dilution of ECL anti-mouse IgG, HRP conjugate (GE Healthcare, Little Chalfont, United Kingdom) for 1 hour at room temperature. Pierce ECL Western Blotting Substrate (Thermo Scientific, Waltham, MA) was used according to manufacturer’s instructions for visualization and the blots were subsequently exposed to x-ray film and developed on a Konica Minolta SRX-101A developer.

Protein gels were stained using Coomassie Brilliant Blue R-250 (BioRad, Hercules, CA) according to manufacturer’s specifications (http://www.bio-rad.com/en-us/product/coomassie-stains).

Protein concentration was determined using the DC Protein Assay kit (BioRad, Hercules, CA) microplate protocol (http://www.bio-rad.com/LifeScience/pdf/Bulletin_9005.pdf).

## RESULTS

### Yeast Pgk1 Fused With the CelTag Binds Efficiently to Cellulose

To test whether a protein fused with the CelTag peptide sequences on its C-terminal end could efficiently bind to cellulose, we chromosomally tagged the ORF for Pgk1, a soluble cytoplasmic protein, as a proof of concept. The CelTag DNA sequences were chromosomally-integrated onto the C-terminus of the *PGK1* gene using homologous recombination (Figure 1A), allowing purification of tagged protein using cellulose as the affinity matrix, eluting from the matrix with TEV protease or SDS (Figure 1B). Previously it was shown that commercially available microgranular cellulose powder has suboptimal accessible surface area and binding capacity compared to regenerated amorphous cellulose (RAC) made by phosphoric acid treatment of the microgranular material (Hong *et al.* 2007; Hong *et al.* 2008; Wang and Hong 2014). Although this treatment causes the cellulose to form more gel-like pellets and is somewhat burdensome for regular use, we compared whether microgranular cellulose or RAC more efficiently pulled down Pgk1-CelTag to a degree that would justify its use. Pgk1-CelTag was pulled down from 450 μL of extract with varying amounts of both regenerated amorphous cellulose and untreated microgranular cellulose. Once the binding was complete the cellulose pellets were eluted in 50 μL of SDS-Page Loading Buffer and subsequently run on a protein gel (Figure 2A). The results confirm that the RAC is more efficient at pulling down the tagged protein, though the microgranular cellulose is at least as efficient as the RAC at recovering tagged protein (76 percent from RAC versus 81 percent from microgranular at the highest concentrations used, Figure 2A). This obviates the need for RAC, since it is more difficult to use, and thus, the subsequent experiments were done using microgranular cellulose. The elutions that gave the most tagged protein from 450 μL of cell extract were the packed pellet of 25 μL or 50 μL microgranular cellulose. Thus, subsequent experiments used approximately 1/20 volume packed cellulose relative to extract volume. Tagged protein was not detected in the unbound fractions (data not shown) suggesting some protein might be unrecoverable from the cellulose matrix.

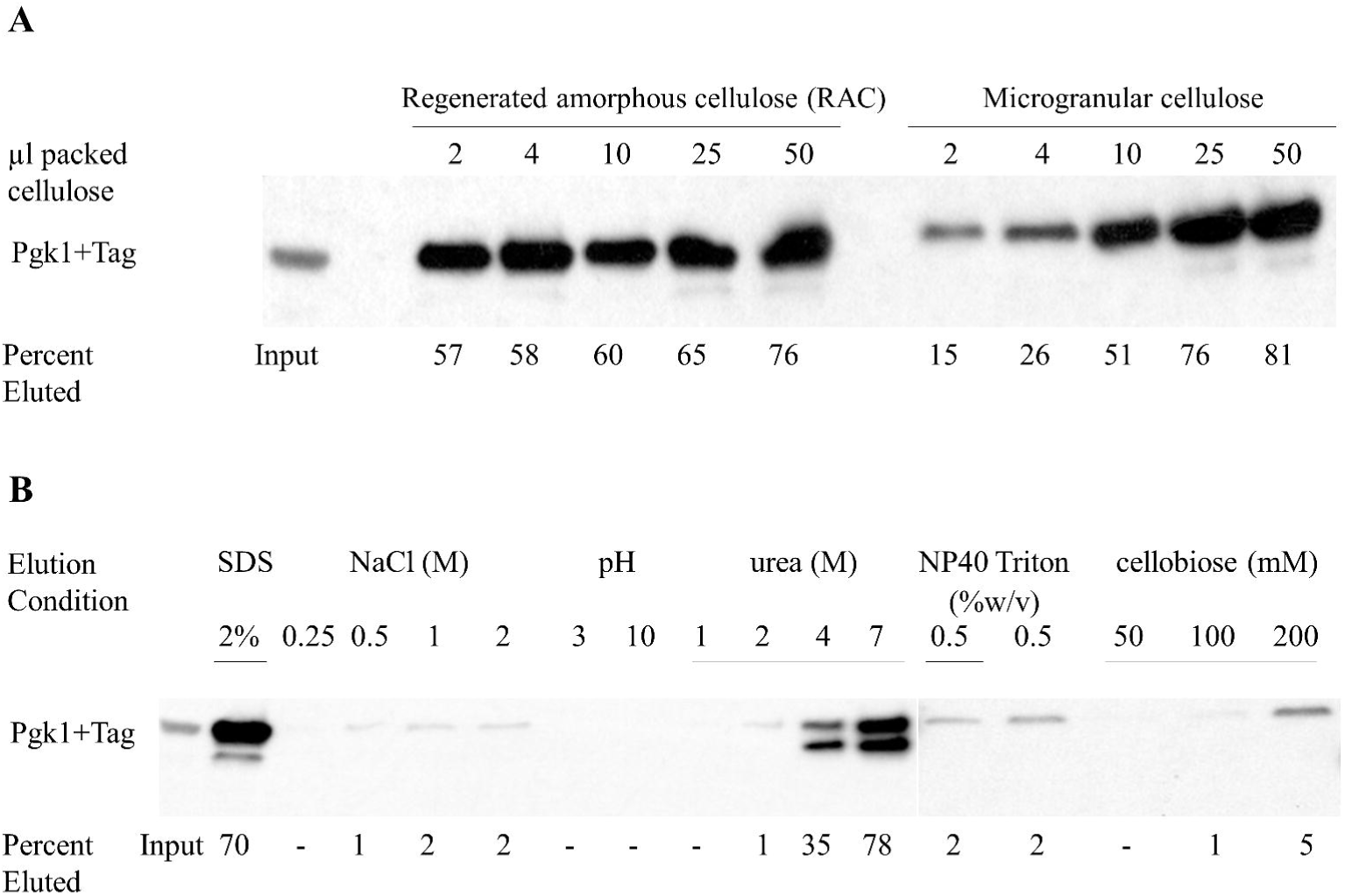

### Pgk1-CelTag Bound to Cellulose is Resistant to Stringent Wash Conditions

Next we sought to determine under what wash conditions the tagged protein would be able to remain bound to the cellulose. Part of the utility of the CelTag could be the high affinity of the CBM3 tag for cellulose under a variety of conditions, allowing stringent washing of the bound protein complexes to remove loosely or non-specifically bound contaminants. Fifty μL of packed microgranular cellulose was used to pull down tagged protein in 450 μL of lysate. Pellets were eluted in 50 μL of binding buffer containing various additions. The CelTag binding seems to be relatively resistant to high salt elutions. Even in the presence of 2 M NaCl, only about 2% of the tagged protein in the input is eluted, indicating that a majority of the tagged protein remains bound to the cellulose (Figure 2B). The tagged protein bound to cellulose is also resistant to elutions in up to 2 M urea with only about 1% of Pgk1-CelTag eluting, compared to 4M urea where about 35% is eluted (Figure 2B). Moderate acidic and basic conditions (pH 3 or 10) also eluted only small amounts of protein (Figure 2B) and binding is almost completely resistant to concentrations of the nonionic detergents NP-40 and Triton X-100 above their critical micelle concentrations, with only 2% of the tagged protein eluted in each case (Figure 2B). Lastly, we investigated whether cellobiose, the cellulose sugar dimer, is able to competitively elute the protein from the matrix. At concentrations just below saturation, cellobiose is only able to elute 5% of the total protein off the cellulose over the time course used (Figure 2B). These data suggest that stringent washing prior to elution could be used to remove the vast majority of molecules that might bind non-specifically to the cellulose matrix or the tagged protein.

### TEV Protease Efficiently and Specifically Elutes Pgk1-CelTag from Cellulose

We next determined the approximate amount of TEV protease needed for optimal release of tagged protein from the cellulose. At the highest concentration of TEV protease, 73% of the tagged protein relative to input is eluted from the cellulose (indicated by a smaller band representing the Pgk1 with the cellulose binding module removed) (Figure 3A, lanes 1 and 3). The TEV elution is also very specific and eliminates most residual non-specific binding contamination seen if the pellet is eluted in SDS (Figure 3B, lanes 2 and 3). To remove the recombinant His6-tagged TEV protease from the elution, we passed the eluted sample through nickel-coated beads (Figure 3B, lane 3 and 4). The total yield and fold-purification using the above method to purify a CelTagged protein is summarized in Figure 3C. In our hands the CelTag purification scheme is able to provide ∼ 470-fold purification of the protein of interest, more importantly recovering about 73% of the total starting protein, using only a single affinity step followed by removal of the TEV protease.

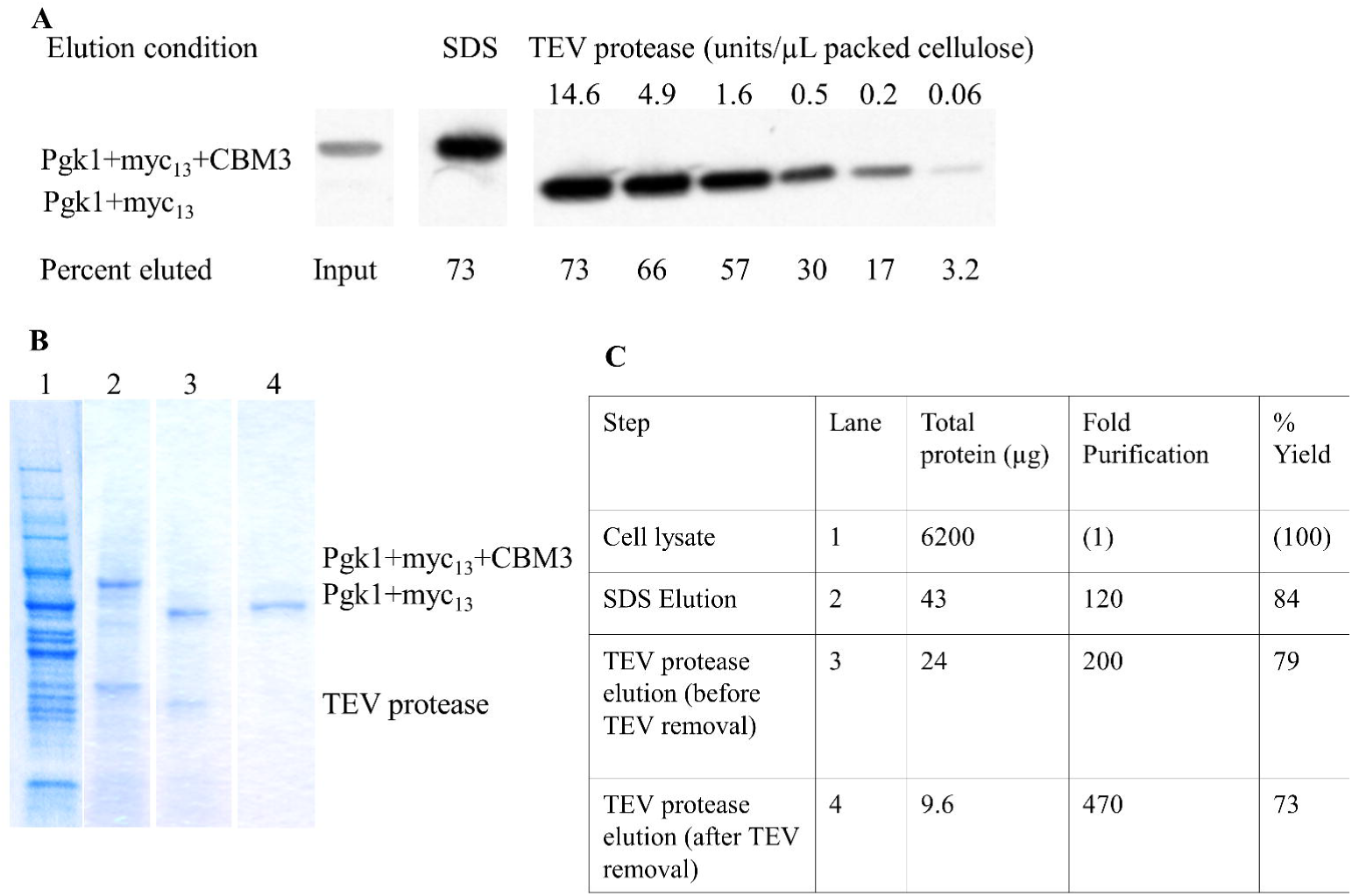

## DISCUSSION

We have shown as a proof of principle that the CelTag affinity tag is able to efficiently recover a majority of total endogenous, tagged protein from crude cellular extracts with nearly complete purity. In contrast to previous uses of the cellulose tags, it appears that using commercially available microgranular cellulose is as effective as using the more labor-intensive RAC at binding the CBM3 domain as an affinity tag. This is useful since microgranular cellulose is much easier to transfer and does not require acid treatment. The recombinant DNA construct encoding the CelTag has been constructed to also contain a 13x c-Myc epitope tag, so that the fragment to be used for tagging a chromosomal ORF can be lifted by PCR to either contain or delete the c-Myc portion as an additional isolation or detection tool.

Another useful aspect of the CelTag is that relatively stringent washing conditions are possible once the tagged proteins are bound to the cellulose support. High salt, non-ionic detergents and urea were all tested and showed that reasonably severe, non-denaturing conditions can be used to minimize non-specific or loosely associated macromolecules from the isolated target. Cellobiose elutions were attempted to see whether the tag could be competed off the cellulose without the use of TEV protease, but with soluble concentrations of cellobiose the tag is unable to be competed off within the tested time frame, likely due to the higher affinity and specificity that the CelTag has for cellulose. Therefore, TEV protease was optimized for efficient elution of CelTagged protein off cellulose.

We envision this protocol to be used for the specific tagging of the chromosomal open reading frames, and purification of rare multipolypeptide, protein-RNA and protein-DNA complexes.

